# A locomotor neural circuit persists and functions similarly in larvae and adult *Drosophila*

**DOI:** 10.1101/2021.04.27.441684

**Authors:** Kristen M. Lee, Chris Q. Doe

## Abstract

Individual neurons can undergo drastic structural changes, known as neuronal remodeling or structural plasticity. One example of this is in response to hormones, such as during puberty in mammals or metamorphosis in insects. However, in each of these examples it remains unclear whether the remodeled neuron resumes prior patterns of connectivity, and if so, whether the persistent circuits drive similar behaviors. Here, we utilize a well-characterized neural circuit in the *Drosophila* larva: the Moonwalking Descending Neuron (MDN) circuit. We previously showed that larval MDN induces backward crawling, and synapses onto the Pair1 interneuron to inhibit forward crawling (Carreira-Rosario et al., 2018). MDN is remodeled during metamorphosis and regulates backward walking in the adult fly. We investigated whether Pair1 is remodeled during metamorphosis and functions within the MDN circuit during adulthood. We assayed morphology and molecular markers to demonstrate that Pair1 is remodeled during metamorphosis and persists in the adult fly. In the adult, optogenetic activation of Pair1 resulted in arrest of forward locomotion, similar to what is observed in larvae. MDN and Pair1 are also synaptic partners in the adult, showing that the MDN-Pair1 interneuron circuit is retained in the adult following hormone-driven pupal remodeling. Thus, the MDN-Pair1 neurons are an interneuronal circuit – i.e. a pair of synaptically connected interneurons – that persists through metamorphosis, taking on new input/output neurons, yet generating similar locomotor behavior at both stages.

## Introduction

Large-scale changes in neuronal morphology and function occur during mammalian puberty (Barendse et al., 2018; Mills et al., 2016; Sisk and Zehr, 2005), as well as several neurobiological disorders including depression (Patel et al., 2019), or chronic pain (Kuner and Flor, 2017). Similarly, major changes in neuronal numbers and type occur as a result of insect metamorphosis (Kanamori et al., 2015; Truman and Reiss, 1976; Yaniv and Schuldiner, 2016). Despite these changes, there are documented cases of individual insect neurons persisting from larval to adult stages. In *Drosophila*, individual motor and sensory neurons have been shown to persist throughout metamorphosis and undergo dramatic remodeling (Consoulas et al., 2002, 2000; Yaniv and Schuldiner, 2016; Yu and Schuldiner, 2014). Similar findings have been reported for the insect mushroom body, where Kenyon cells partners (projection neurons, DANs) exist at both larval and adult stages (Li et al., 2020; Marin et al., 2005). Yet, it remains unclear whether the remodeled neurons re-establish connectivity with the identical neurons in the larva and adult.

During *Drosophila* metamorphosis the animal changes from a crawling limbless larva to a walking six-legged adult (Riddiford, 1980; Riddiford et al., 2003). Despite the obvious differences, some behaviors are similar: both larvae and adults undergo forward locomotion in search of food, backward locomotion in response to noxious stimuli, and pausing in between antagonistic behaviors (Carreira-Rosario et al., 2018). We and others identified a neuron that, when activated, can trigger backward locomotion in both larvae and adults (Bidaye et al., 2014; Carreira-Rosario et al., 2018; Sen et al., 2017), despite the obvious differences in limbless and six-legged locomotion. This neuron, named Moonwalker/Mooncrawler Descending Neuron (MDN) is present in two bilateral pairs per brain lobe, with all four MDNs having similar synaptic partners, and all four MDNs capable of eliciting backward larval locomotion in larvae (Carreira-Rosario et al., 2018). Larval MDNs induce backward locomotion via the coordinate arrest of forward locomotion followed by the initiation of backward locomotion. Halting forward locomotion is done via activation of the Pair1 descending interneuron, which inhibits the A27h premotor neuron, to prevent it from inducing forward locomotion (Carreira-Rosario et al., 2018). Activating backward locomotion is likely to be due, in part, to MDN activation of the A18b premotor neuron, which is specifically active during backward locomotion (Carreira-Rosario et al., 2018). Thus, MDN-Pair1 are synaptically coupled members of a locomotor circuit in the *Drosophila* larva.

Here we follow our previous work showing that MDN is remodeled during metamorphosis and persists into the adult (Carreira-Rosario et al., 2018) by asking: Is the MDN partner neuron Pair1 also maintained in the adult? Does the adult Pair1 induce an inhibition in forward locomotion, similar to its role in larvae? And, are the adult Pair1 and MDN synaptically coupled? We find that all of these questions are answered in the affirmative, showing that the core MDN-Pair1 decision-making circuit (a pair of synaptically-connected interneurons) persists from larva to adult, despite profound remodeling during metamorphosis, and that this circuit coordinates forward/backward locomotion in both larvae and adults.

## Results

### The Pair1 neuron persists from larval to adult stages

To determine if Pair1 neurons were present in the adult, we mapped expression of a Pair1-Gal4 line (*R75C02-Gal4*) from early larval to adult stages. We identified the larval Pair1 neurons based on their characteristic cell body position in the medial subesophageal zone (SEZ), dense local ipsilateral dendritic arborizations (defined as dendritic based on enrichment for post-synapses in the TEM reconstruction of the larval Pair1 neuron; Figure 1 – supplement 1), and contralateral axons descending into the ventral nerve cord (VNC) in an extremely lateral axon tract (Carreira-Rosario et al., 2018). Using the Pair1-Gal4 line, we could identify Pair1 neurons with this morphology at 28h and 96h after larval hatching (ALH; Figure 1A,B). The Pair1 neuron cell bodies and proximal neurites could still be observed at 24 hrs after pupal formation (APF), but virtually all of the dendridic processes and descending axonal process had been pruned (Figure 1C, only one neuron labeled). This is expected, given that many or all neurons undergo axon/dendrite remodeling during metamorphosis (Kanamori et al., 2015; Truman and Reiss, 1976; Yaniv and Schuldiner, 2016). At 48 hrs APF, Pair1 neurons exhibited dendritic branching in the SEZ and a descending axon into the VNC, regaining morphological features similar to that of larval Pair1 neurons (Figure 1D). The axon innervated the T1 (prothoracic) neuropil and descended further down the VNC. These morphological features were maintained into the adult fly, where we could trace the Pair1 axon to primarily innervate the T1 neuropil (Figure 1E), with less extensive innervation of the mesothoracic (T2) and metathoracic (T3) neuropils.

**Figure 1.**
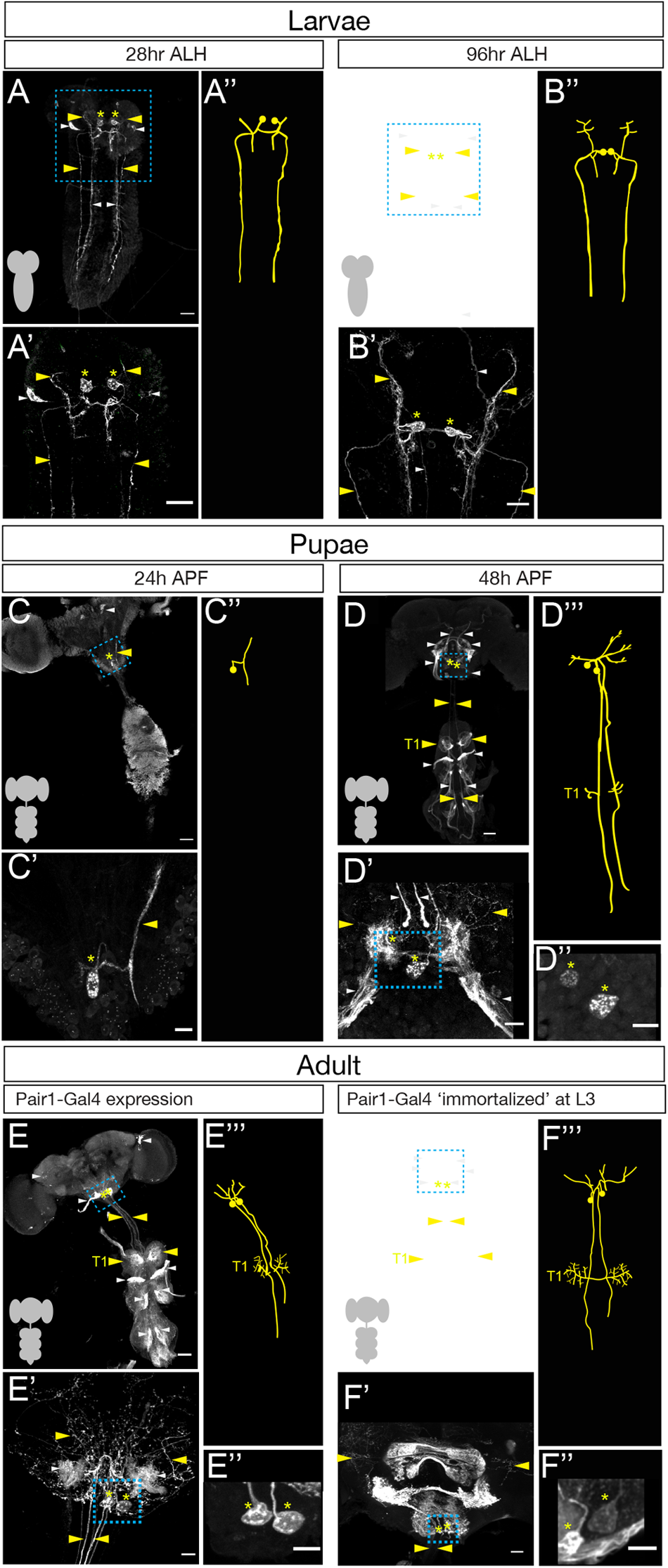
The Pair1 neuron persists from larval to adult stages. (**A-B**) Pair1 neurons (cell body: yellow asterisk; neurites: yellow arrowhead) in the larval CNS (gray outline) at 28h ALH (A) and 96h ALH (B). Here and in subsequent panels are maximum intensity projections of confocal sections containing the Pair1 neurons; anterior, up; dorsal view. Significant ‘off-target’ expression marked with white arrowheads. Scale bar, 50 μm. (**A’-B’**) Enlargement of the brain regions boxed in A,B. Scale bar, 20 μm. (**A’’-B’’**) Tracing to show Pair1 neuron morphology. Genotype: *+; UAS-myr::GFP; R75C02-Gal4*. (**C-D**) Pair1 neurons (cell body: yellow asterisk; neurites: yellow arrowhead) in the pupal CNS (gray outline) at 24h APF (C) and 96h APF (D). Significant ‘off-target’ expression marked with white arrowheads. Scale bar, 50 μm. (**C’-D’**) Enlargement of the brain regions boxed in C, D; cell body: yellow asterisk, neurites: yellow arrowhead. Scale bar, 10 μm. (**C’’**) Tracing to show Pair1 neuron morphology. (**D’’**) Focal plane showing Pair1 cell bodies (region boxed in D’, cell body marked with yellow asterisks). Scale bar, 10 μm. (**D’’’**) Tracing to show Pair1 neuron morphology. Note that Pair1 can be followed to T1 in the 3D confocal stack but is difficult to represent here due to fasciculation of Pair1 with off-target neurons. Genotype: *+; UAS-myr::GFP; R75C02-Gal4*. (**E**) Pair1 neurons (cell body: yellow asterisk; neurites: yellow arrowhead) in the 4d adult CNS (gray outline) Significant ‘off-target’ expression marked with white arrowheads. Scale bar, 50 μm. (**E’**) Enlargement of the brain region boxed in E. Scale bar, 10 um. (**E’’**) Focal plane showing Pair1 cell bodies (region boxed in E’, cell body marked with yellow asterisks). Scale bar, 10 μm. (**E’’’**) Tracing to show Pair1 neuron morphology. Genotype: *+; UAS-myr::GFP; R75C02-Gal4*. (**F**) Pair1 neurons (cell body: yellow asterisk; neurites: yellow arrowhead) permanently labeled at 96h ALH and visualized in the 4d old adult. See methods for details. Significant ‘off-target’ expression marked with white arrowheads. Scale bar, 50 μm. (**F’**) Enlargement of the brain region boxed in F; Pair1 cell body: yellow asterisk; Pair1 neurites: yellow arrowhead. Scale bar, 10 μm. (**F’’**) Focal plane showing Pair1 cell bodies (region boxed in F’, cell body marked with yellow asterisks). Scale bar, 10 μm. (**F’’’**) Tracing to show Pair1 neuron morphology. Genotype: *Hs-KD,3xUAS-FLP; 13xLexAop(KDRT*.*Stop)myr:smGdP-Flag/+; 13xLexAop(KDRT*.*Stop)myr:smGdP-V5, 13xLexAop(KDRT*.*Stop)myr:smGdP-HA, nSyb(FRT*.*Stop)LexA::p65*

Although we can use the Pair1-Gal4 line to track neurons with Pair1 morphological features from larva to adult, it remains possible that the Gal4 line switches off in Pair1 and switches on in a similar descending neuron at a stage in between those we assayed. To conclusively demonstrate that the larval Pair1 neuron survives into adulthood, we used a genetic technique to permanently label or “immortalize” the larval Pair1 neurons and assay for their presence in the adult brain. Briefly, the method acheives spatial specificity by using Pair1-Gal4 to drive UAS-FLP which removes a stop cassette from nSyb-FRT-stop-FRT-LexA resulting in permanent LexA expression in Pair1-Gal4 neurons; it acheives temporal specificity (e.g. labeling only larval Pair1-Gal4+ neurons) by using a heat inducible KD recombinase to “open” the lexAop-KDRTstopKDRT-HA reporter (see Methods for additional details). Thus, a heat shock will permanently label all Pair1-Gal4+ neurons at the time of heat shock. We immortalized Pair1 neurons in the larva, and assayed expression in the adult, and observed the two bilateral Pair1 neurons, based on characteristic medial SEZ cell body position, local ipsilateral arbors and contralateral descending axons that preferentially innervate the prothoracic neuropil (Figure 1F). Pair1 innervation is clearer in neurons immortalized during larval stages, which reduces the off-target neuron expression in the adult VNC, and reveals an greatly enriched level of innervation in the T1 neuropil (Figure 1F).

The Pair1-Gal4 line is expressed in several off-target neurons in addition to Pair1. One of these, a sensory neuron from the proboscis, can be reduced from the adult Pair1 pattern by removing the proboscis a day prior to analysis (see Methods) but is present at the 48 hr APF timepoint (Figure 1D, E). In addition, there are off-target neurons that innervate all three thoracic neuropils (T1-T3), obscuring Pair1 innervation (Figure 1E). We took advantage of the sparse labeling of the immortalization genetics and found brains that maintained preferential targeting of Pair1 to the prothoracic neuropil but lacked T1-T3 off-target innervation, confirming that they are indeed off-target neurons (Figure 1F-F’’’). We conclude that the Pair1 neurons are present from larval to adult stages, and that the Pair1 neurons are enriched for postsynaptic partners in the T1 neuromere.

### Pair1 neurons maintain the same molecular profile from larval to adult stages

If Pair1 neurons persist from larva to adult, they may express the same transcription factor (TF) profile at both stages. We screened a small collection of TF markers for expression in the larval and adult Pair1 neurons, and in all cases we found identical expression (Figure 2). Larval and adult Pair1 neurons expressed Hunchback (Hb), Sex combs reduced (Scr), and Bicoid (Bcd); but did not express Visual system homeobox (Vsx1) or Nab (Figure 2). These results support the conclusion that Pair1 persists from larva to adult, maintaining both molecular and morphological features, and raises the interesting possibility that the three TFs (Hb, Bcd, Scr) may provide a molecular code that directs both larval and adult Pair1 morphology and/or connectivity (see Discussion).

**Figure 2.**
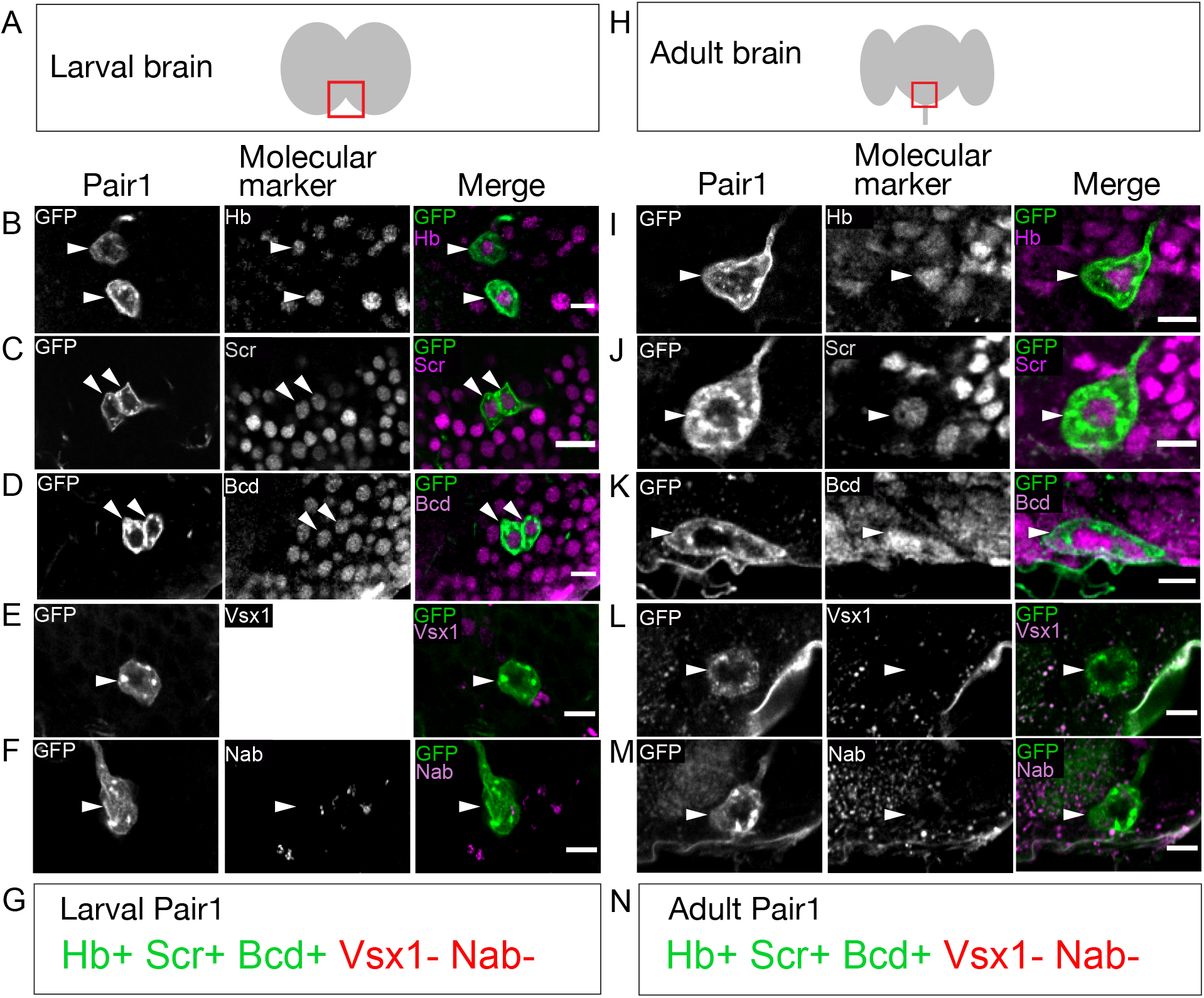
The Pair1 neuron expresses the same molecular markers at larval and adult stages. (**A**) Schematic of the larval brain showing region of Pair1 neurons (red box) enlarged in panels below. Anterior up, dorsal view. (**B-G**) Larval Pair1 neurons (left column), indicated markers (middle column), and merge (right column) at 28h ALH. In some cases the second Pair1 neuron is out of the focal plane, but both Pair1 neurons always have the same gene expression profile. Markers detect the following transcription factors: Hb, Hunchback; Scr, Sex combs reduced; Bcd, Bicoid; Vsx1, Visual system homeobox 1; and Nab. Scale bar, 5 μm. (G) Summary: marker expression matches that in adults. Genotype: *+; UAS-myr::GFP; R75C02-Gal4*. (**H**) Schematic of the adult brain showing region of Pair1 neurons (red box) enlarged in panels below. Anterior up, dorsal view. (**I-N**) Adult Pair1 neurons (left column), indicated markers (middle column), and merge (right column) in 4d old adult. Scale bar, 5 μm. (N) Summary: marker expression matches that in larvae. Genotype: *+; UAS-myr::GFP; R75C02-Gal4*.

### Pair1 activation arrests forward locomotion in adults

We previously showed that larval MDN persists in adults and can induce backward locomotion at both stages despite the obvious difference in motor output – limbless crawling vs. six-legged walking (Carreira-Rosario et al., 2018). This raised the question of whether the adult Pair1 neuron also maintains its larval function, i.e. to pause forward locomotion. To test this hypothesis, we used Pair1-Gal4 to express the red light-gated cation channel CsChrimson (Chrimson) to activate Pair1 neurons in the adult. Experimental flies were fed all-*trans* retinal (ATR; required for Chrimson function) whereas control flies were fed vehicle only.

Control flies exposed to red light did not pause or arrest forward locomotion, did not show an increased probability of pausing, and did not have a decrease in distance traveled during the stimulus interval. In contrast, experimental flies expressing Chrimson in Pair1 neurons showed a near complete arrest of forward locomotion, an increased probability of pausing, and a reduced distance traveled during the stimulus interval (Figure 3A-C; Figure 3-Supplement 1). These effects were reversed after turning off the red light, with the exception of a slightly reduced distance travelled, likely due to a lingering physiological effect of the 30 sec Pair1 activation (Figure 3A,C). Pair1 activation resulted in an increase in immobile flies (Figure 3E) and a corresponding decrease in whole body translocation (defined as “large movements”, Figure 3F). Importantly, Pair1 activation did not prevent small body part movements such as those involved in grooming (defined as “small movements”, Figure 3G). Note that Pair1-Gal4 off-target expression is common but variable from fly to fly, whereas its expression in Pair1 neurons is fully penetrant; because the Chrimson-induced behavior is also fully penetrant, we conclude that the arrest in forward locomotion is due to Chrimson activation of the Pair1 neurons. We conclude that Pair1 activation prevents a single behavior – forward locomotion – but does not produce general paralysis or interfere with non-translocating limb movements.

**Figure 3.**
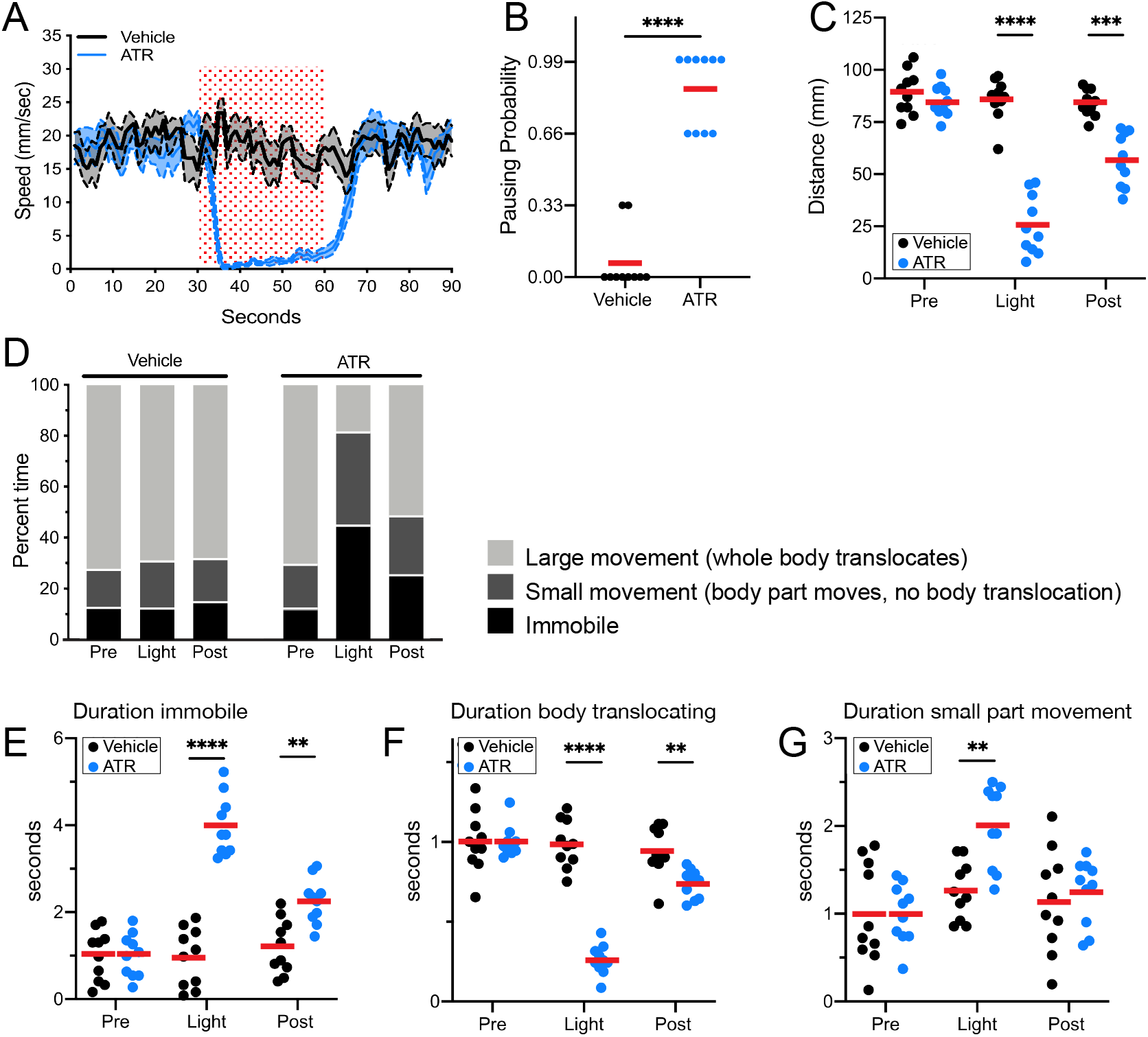
Pair1 activation for 30s arrests forward locomotion but does not cause paralysis in adults. (**A**) Speed (mm/sec) of adult flies expressing Chrimson in Pair1 neurons following neuronal activation (+ATR, blue) or no activation (vehicle control, black) in a closed loop arena. Speed was recorded for the 30s prior to activation, the 30s light-induced activation (red stipple), and 30s after activation. Mean ± S.E.M, n = 10. Genotype for this and all subsequent panels: *UAS-CsChrimson::mVenus; +; R75C02-Gal4*. (**B**) Probability of forward locomotion pausing upon light-induced Pair1 activation (ATR treatment, blue) compared to vehicle control (black). Statistics: t-test, p < 0.001; n = 10. (**C**) Total distance traveled pre-light stimulus (“pre”), during the light stimulus (“light”) and post-light stimulus (“post”) (terminology used here and in subsequent panels) of flies fed ATR (Pair1 activation, blue) compared to controls (fed vehicle, no Pair1 activation, black). Statistics: two-way ANOVA: drug treatment, F(1, 18) = 111.3, p < 0.0001; time, F(1.867, 33.61) = 47.03, p < 0.0001; interaction F(2, 26) = 38.24, p < 0.001; Bonferroni’s multiple comparisons between drug treatments within each timepoint: pre, p > 0.9999; light, p < 0.0001; post, p = 0.0001; n = 10. (**D**) Percent time doing large movements (whole body translocation, light grey), small movements (body part movement but no translocation, dark grey) or no movements (immobile, black) of flies fed vehicle (left side) or ATR (right side) during each time phase (pre, light, post). (**E**) Normalized duration of time spent <u>immobile</u> during each timepoint (pre, light, post) for flies fed ATR (Pair1 activation, blue) compared to controls fed vehicle (black). Statistics: two-way ANOVA: drug treatment, F(1, 18) = 112.8, p < 0.0001; time, F(1.930, 34.74) = 25.55, p < 0.0001; interaction, F(2, 36) = 27.81, p < 0.0001; Bonferroni’s multiple comparisons between drug treatments within each timepoint: pre, p > 0.9999; light, p < 0.0001; post, p = 0.0022; n = 10. (**F**) Normalized duration of time spent doing <u>small movements</u> during each timepoint (pre, light, post) for flies fed ATR (Pair1 activation, blue) compared to controls fed vehicle (black). Statistics: two-way ANOVA: drug treatment, F(1, 18) = 5.111, p = 0.036; time, F(1.923, 34.62) = 10.82, p = 0.0003; interaction, F(2, 36) = 4.225, p = 0.0225; Bonferroni’s multiple comparisons between drug treatments within each timepoint: pre, p > 0.9999; light, p = 0.0022; post, p > 0.9999; n = 10. (**G**) Normalized duration of time spent doing <u>large movements</u> during each time phase (pre, light, post) for flies fed ATR (Pair1 activation, blue) compared to controls fed vehicle (black). Statistics: two-way ANOVE: drug treatment, F(1, 18) = 53.56, p < 0.0001; time, F(1.869, 33.64) = 53.44, p < 0.0001; interaction, F(2, 36) = 52.20, p < 0.0001; Bonferroni’s multiple comparisons between drug treatments within each timepoint: pre, p > 0.9999; light, p < 0.0001; post, p = 0.0074; n = 10.

### MDN and Pair1 are synaptic partners during adulthood

Given that MDN and Pair1 are synaptic partners in the larvae (Figure 1-Supplement 1), MDN and Pair1 persist into adulthood (Figure 1 and 2), and MDN and Pair1 both regulate the same behavior in larvae and adults (Figure 3) (Carreira-Rosario et al., 2018), we hypothesized that MDN and Pair1 may also be synaptic partners during adulthood. To test this hypothesis, we used the MDN-LexA and Pair1-Gal4 to label MDN and Pair1 neurons individually in the same animal (Figure 4A,B). We observed MDN and Pair1 neurites in close proximity to each other (Figure 4C-E). To determine if MDN and Pair1 are synaptic partners in this region of neuropil, we utilized t-GRASP (targeted GFP reconstitution across synaptic partners), an activity-independent method to label synaptic contact sites (Shearin et al., 2018). Control flies only expressing pre-t-GRASP in MDN did not have detectable t-GRASP signal (Figure 4F). However, flies expressing pre-t-GRASP in MDN and post-t-GRASP in Pair1 had t-GRASP signal, indicating that MDN and Pair1 form synapses (Figure 4G). We conclude that MDN and Pair1 are synaptic partners during adulthood.

**Figure 4.**
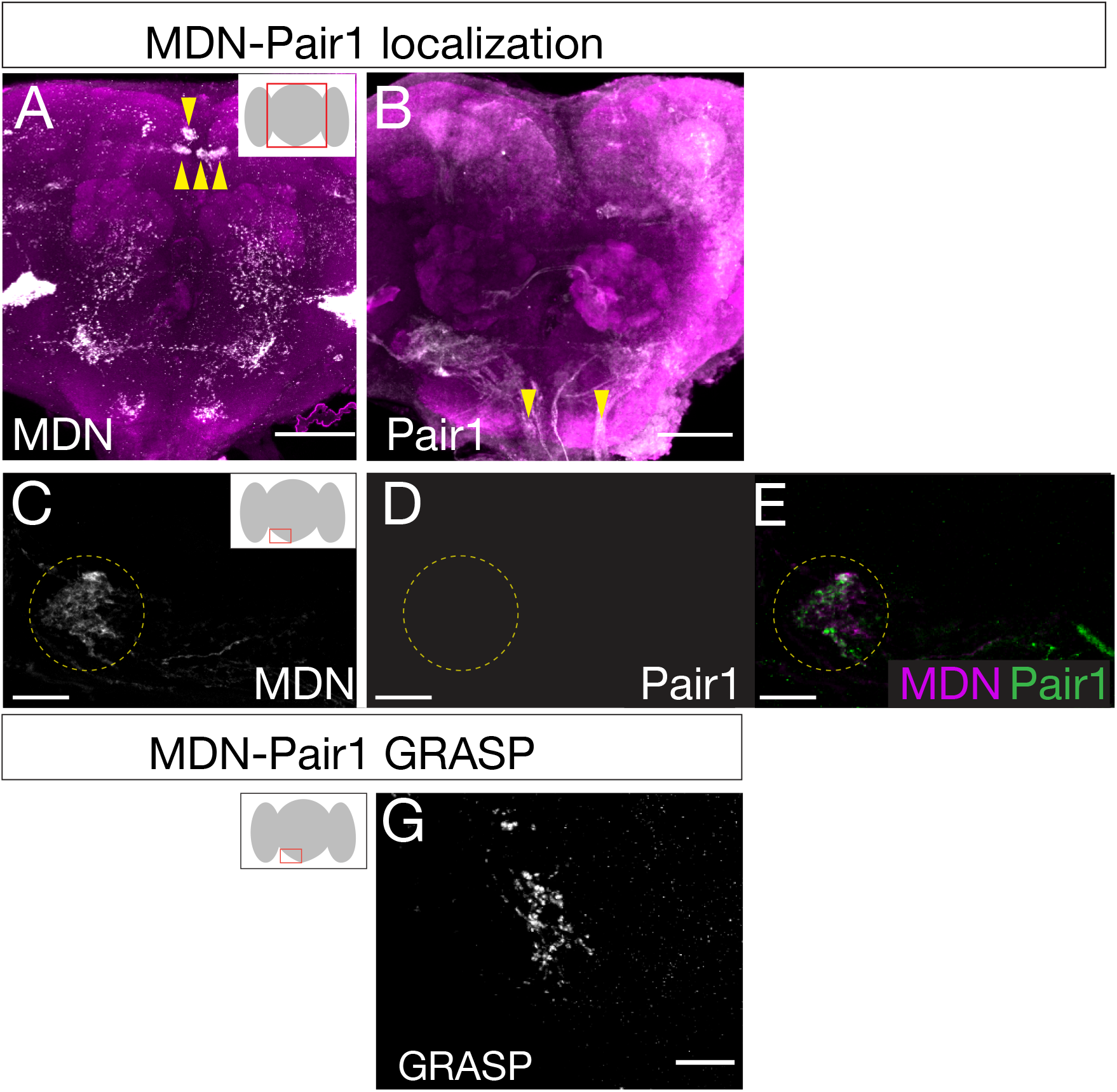
MDN and Pair1 are synaptic partners in adults. (**A, B**) MDN neurons (A) and Pair1 neurons (B) in the adult central brain. Neurons are in white, nc82 counterstain in magenta for whole brain orientation; cell bodies marked by yellow arrowheads. Here and in subsequent panels shows maximum intensity projection of volume; anterior, up; dorsal view. Scale bar, 50 μm. Genotype: *UAS-mCD8::RFP, LexAop-mCD8::GFP; VT044845-LexA; R75C02-Gal4* (**C-E**) MDN neurites (C), Pair1 neurites (D), and merge (E) in the subesophageal ganglion (red box in schematic). Scale bar, 10 μm. Genotype: *UAS-mCD8::RFP, LexAop-mCD8::GFP; VT044845-LexA; R75C02-Gal4*. (**F**) No detectable t-GRASP signal was observed in the subesophageal ganglion without expression of the pre-t-GRASP fragment in MDN. Scale bar, 10 μm. Genotype:;; *LexAop-pre-t-GRASP, UAS-post-t-GRASP/R75C02-Gal4*. (**G**) t-GRASP signals between MDN and Pair1 were observed in the subesophegeal ganglion. Scale bar, 10 μm. Genotype:; *VT044845-LexA*; *LexAop-pre-t-GRASP, UAS-post-t-GRASP/R75C02-Gal4*.

## Discussion

Together with our earlier work (Carreira-Rosario et al., 2018), our results here show that a core decision-making circuit is preserved from larval stages into the adult. This decision-making circuit contains MDN and its monosynaptically-coupled Pair1 neuron, allowing the fly to switch between antagonistic behaviors: forward versus backward locomotion. Our work raises several interesting questions: Do many other larval neural circuits persist and have similar function in adults? Are the cues that establish MDN-Pair1 connectivity in the larvae also used to re-establish MDN-Pair1 connectivity in the adult?

How much of the larval MDN-Pair1 circuit is maintained into the adult? The larval circuit contains the MDN partners Pair1, ThDN, and A18b, and the Pair1 partner A27h (Figure 5). In addition to MDN, we show here that Pair1 is maintained. There is no Gal4 line or markers for the ThDN neuron, and the only A18b line has extensive off-target expression, so the fate of these two neurons is unknown. In contrast, the A27h interneuron, which regulates forward crawling in the larvae, undergoes apoptosis during pupal stages (data not shown) and thus does not regulate forward walking in the adult. This is not surprising as the A27h neurons are located in the abdominal segments, which do not have a role in adult walking.

**Figure 5.**
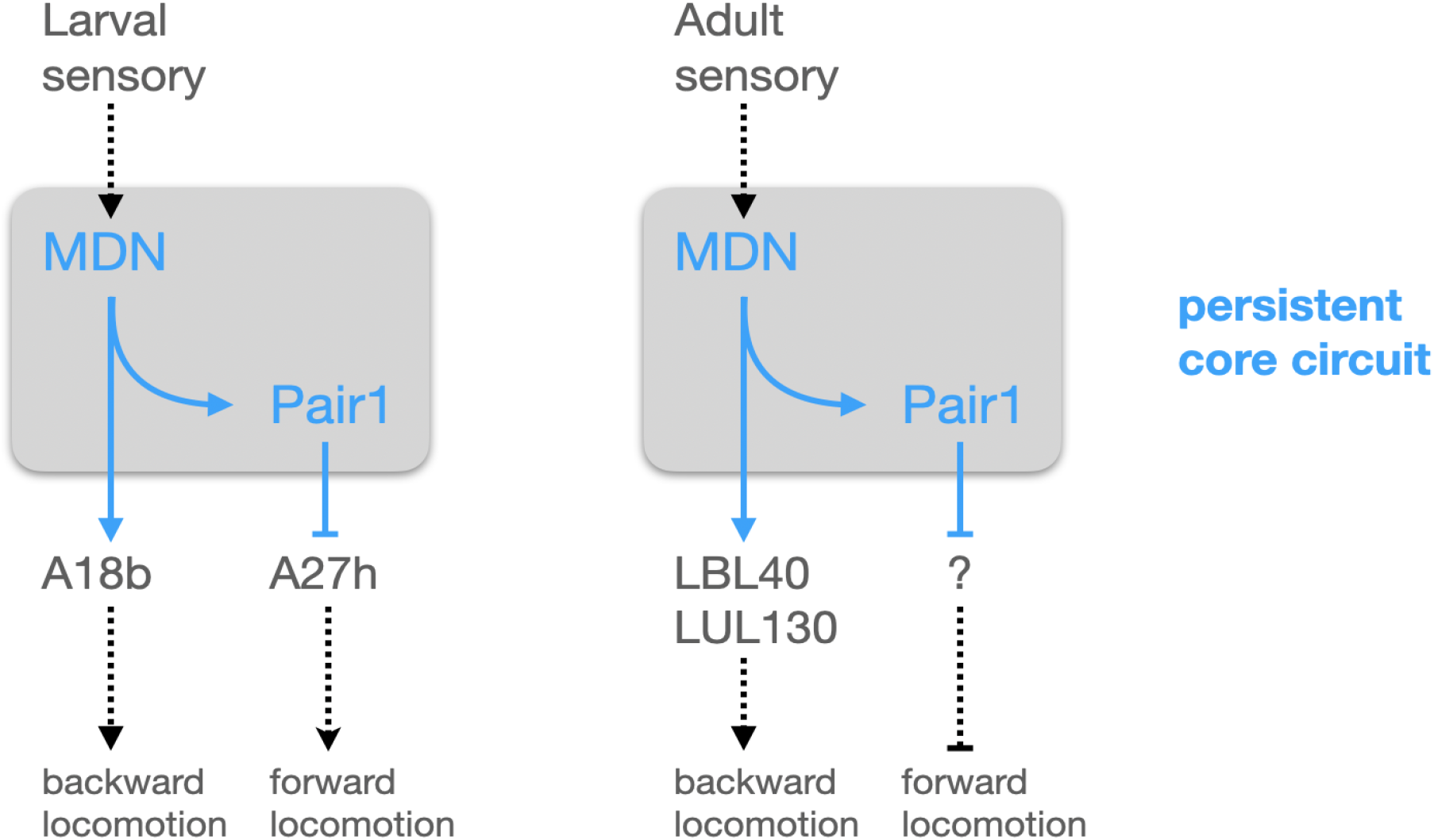
Model describing the MDN-Pair1 circuit in larval and adult stages. In the larva, activation of MDN by upstream sensory neurons activates backward crawling and inhibits forward crawling. MDN neurons synapse onto A18b and Pair1 interneurons. A18b subsequently regulates backward crawling by synapsing onto motor neurons. Pair1 synapses onto and inhibits the pre-motor neuron A27h, which generates forward locomotion when activated. In the adult, activation of MDN by upstream sensory neurons activated backward walking and inhibits forward walking. MDN neurons synapse onto LBL40, LUL120 and Pair1. Activation of LBL40 and LUL130 generates backward walking. Activation of Pair1 generates a pausing behavior, likely through the inhibition of neurons generating forward locomotion. In both larvae and adult, MDN and Pair1 neurons (blue) persist and function as a core circuit to regulate locomotion.

How much of the adult MDN-Pair1 circuit is present in the larvae? Recent work mapping the adult MDN circuit has identified over 30 VNC neurons downstream of MDN, including the LBL40 and LUL130 neurons required for hindleg backward stepping (Figure 5; Feng et al., 2020). Recent work has also identified adult neurons important for forward walking (Bidaye et al., 2020), but their relationship to adult MDN is unknown. In the future, it will be interesting to see if any of these adult neurons are present in the larvae, particularly those regulating forward and backward walking, and determine if they are also MDN or Pair1 target neurons.

Elegant recent work has shown that initiation of forward walking requires the forelegs, innervated by motor neurons in the prothoracic segment, whereas initiation of backward walking requires the hindlegs, innervated by motor neuron in the metathoracic segment (Feng et al., 2020). This is consistent with our finding that the adult Pair1 neuron innervates the prothoracic neuropil, a site well-positioned to arrest motor activity driving foreleg stepping and initiation of forward locomotion. Similarly, adult MDN synaptic partners primarily innervate the metathoracic neuromere (Feng et al., 2020), a good location for inducing hindleg stepping and initiation of backward walking. A similar spatial segregation is likely to occur in the larva, where forward crawling is induced by A27h in posterior segments, and backward crawling is induced in anterior segments(Fushiki et al., 2016; Tastekin et al., 2018).

In larvae, Pair1 activation only causes a pause in forward locomotion – after pausing, the animal returns to baseline speed regardless of the red-light stimulus duration (Carreira-Rosario et al., 2018; Tastekin et al., 2018). In contrast, Pair1 activation in adults results in a persistent arrest in forward locomotion for the duration of the red light stimulus, although non-translocating limb movements are not affected. We speculate these differences may be due to differences in Pair1 downstream synaptic partners, with more redundancy in the larval circuit. Understanding the function of the neurons downstream of adult Pair1 in the T1 neuromere is likely to provide insights into these behavioral differences.

Here we identify a transcription factor combination (Hb, Bcd, Scr) that persists in Pair1 neurons from larvae to adults. It is intriguing to speculate that these transcription factors may be required for cell surface molecule expression used to establish the MDN-Pair1 synaptic specificity in the embryo as well as to re-establish MDN-Pair1 synaptic specificity following pupal remodeling. Perhaps these transcription factors drive expression of the same cell surface molecules at both stages, or even continuously to maintain functional connectivity.

Individual neurons that have similar functions in larva and adults have been identified, including select motor neurons, sensory neurons (Consoulas et al., 2002, 2000; Levine, 1984; Truman, 1992; Weeks, 2003), and Kenyon cells of the mushroom body (Eichler et al., 2017; Li et al., 2020). However, it remains unknown whether any of their synaptic partners also persist and retain the same pattern of connectivity. Our work is the first, to our knowledge, to show that a pair of synaptically connected interneurons – the core of a decision-making circuit – can persist from larva to adult and perform similar functions at both stages. Remarkably, both MDN and Pair1 undergo dramatic pruning and regeneration events during metamorphosis, only to re-form synapses with each other following neuronal remodeling. This suggests that synapse specificity cues are maintained from the late embryo, where MDN-Pair1 connectivity is first established, into pupal stages, where MDN-Pair1 connectivity is re-established. The importance of the MDN-Pair1 decision-making circuit is highlighted by its persistence from embryo to adult, despite adapting to different sensory input and motor output at each stage. Perhaps other descending or ascending interneurons will also persist into adults, switching inputs and outputs as needed. Indeed, the idea that a core decision-making circuit that is stable across developmental stages is supported by recent elegant TEM reconstruction of neural circuits at different stages of *C. elegans* development (Witvliet et al., 2020). Here the authors conclude that “Across maturation, the decision-making circuitry is maintained whereas sensory and motor pathways are substantially remodeled.” These results, together with ours, raise the possibility that preservation of decision-making interneuron circuit motifs may be functional modules that can be used adaptively with different sensorimotor inputs and outputs. The presence of this circuit motif in both flies and worms suggests that it may be an ancient evolutionary mechanism for assembling sensorimotor circuits.

## Materials and Methods

### Fly husbandry

All flies were reared in a 25°C room at 50% relative humidity with a 12 hr light/dark cycle. All comparisons between groups were based on studies with flies grown, handled and tested together.

### Fly Stocks

1. *R75C02-Gal4* (Pair1 line; BDSC #39886)
2. *UAS-myr::GFP* (BDSC #32198)
3. *UAS-CsChrimson::mVenus* (Vivek Jayaraman, Janelia Research Campus)
4. *VT044845-lexA* (adult MDN line; a gift from B. Dickson, Janelia Research Campus)
5. *UAS-mCD8::RFP, LexAop-mCD8::GFP;;* (BDSC #32229)
6. *LexAop-pre-t-GRASP, UAS-post-t-GRASP* (BDSC #79039)
7. *Hs-KD,3xUAS-FLP; 13xLexAop(KDRT*.*Stop)myr:smGdP-Flag/ CyO-YFP*; *13xLexAop(KDRT*.*Stop)myr:smGdP-V5, 13xLexAop(KDRT*.*Stop)myr:smGdP-HA, nSyb-(FRT*.*Stop)-LexA::p65/R75C02-Gal4* (line to permanently label Pair1; Doe Lab; modified from (Ren et al., 2016))

### Gal4 driver “immortalization”

Immortalization flies (see genotype #7, above) were allowed to lay eggs for four hours. Newly hatched larvae were placed in a food vial, and at 96 hours ALH the food vial was partially submerged in a 37°C water bath for 5 minutes, allowing the hs-KD to act as a recombinase to remove the KDRT Stop cassette, resulting in nSyb-LexA driving HA expression permanently in the neurons expressing Pair1-Gal4 at the time of heat shock (96h ALH). After the heat shock, larvae in the food vial recovered at 18°C for 5 minutes, and then grown to adulthood at 25°C.

### Immunostaining and imaging

Standard confocal microscopy and immunocytochemistry methods were performed as previously described (Carreira-Rosario et al., 2018). Primary antibodies used recognize: GFP (rabbit, 1:500, ThermoFisher, Waltham, MA; chicken, 1:1500, Abcam12970, Eugene, OR), HA (rat, 1:100, Sigma, St. Louis, MO), Hunchback (mouse, 1:400, AbcamF18-1G10.2), Sex combs reduced (mouse, 1:10, Developmental Studies Hybridoma Bank, Iowa City, IA), Bicoid (rat, 1:100, John Reinitz, University of Chicago, Illinois), Vsx1 (guinea pig, 1:500, Claude Desplan, NYU, New York), Nab (guinea pig, 1:500, Stefan Thor, University of Queensland, Brisbane, Australia) and t-GRASP signal (rabbit GFP G10362, 1:300, Invitrogen). Secondary antibodies were from Jackson ImmunoResearch (Donkey, 1:400, West Grove, PA). Confocal image stacks were acquired on a Zeiss 800 microscope. All images were processed in Fiji (https://imagej.new/Fiji) and Adobe Illustrator (Adobe, San Jose, CA). Images were processed as described previously (Carreira-Rosario et al., 2018). The primary neurites of Pair1 were traced using the Simple Neurite Tracer in Fiji.

### Adult Behavioral Experiment

Adult behavior was assayed using two arenas, a closed loop arena (Figure 3) and an open field arena (Figure 3 – Supplement 1). For the closed loop arena, adult female flies 1 day after eclosion were transferred to standard cornmeal fly food supplemented with 100 mL 0.5 mM ATR or 100% ethanol for 4 days (changed every 2 days). Animals, with intact wings, were starved for 4 hrs and then placed in arenas and their behavior was recorded as described previously (Carreira-Rosario et al., 2018). Flies were exposed to low transmitted light, red light, and low transmitted light again for 30 sec each. This was done three times for each animal. To calculate different parameters, the recorded videos were tracked and analyzed using the CalTech Fly Tracker (Fontaine et al., 2009) and JABA (Kabra et al., 2013). The speed, distance and behavior reported were specific to the first trial. The reported speeds are the average speed of each second. The pausing probability was calculated as previously described (Carreira-Rosario et al., 2018). “Pre” defines the 30 secs prior to red light exposure, “light” defines the 30 secs of red light exposure and “post” defines the 30 secs after red light exposure. Immobile movements were defined as the fly not translocating and not moving other body parts. Small movements were defined as the fly not translocating but moving body parts (i.e. grooming, moving wings). Large movements were defined as the fly translocating its body. All behavior measures were normalized by dividing them by the group average “pre” values.

For the open field arena, adult flies were fed ATR and vehicle as described above. 3 animals were placed in a circular arena with a diameter of 14.5 cm and height of 0.5 cm. After 5 min for environmental acclimation, animal behavior was recorded at 25 FPS using a Basler acA2040-25gm GigE camera under infrared light for 4 sec followed by 4 secs under red light and another 4 sec under infrared light, as described previously (Risse et al., 2013). The was repeated 3 times, and tracked and analyzed as described above.

### Statistics

All statistical analysis (t-test, one-way and two-way ANOVA with Bonferroni’s multiple comparison tests) were performed with Prism 9 (GraphPad software, San Diego, CA). Numerical data in graphs show individual measurements (animals), means (represented by red bars) or means ± S.E.M. (dashed lined), when appropriate. The number of replicates (n) is indicated for each data set in the corresponding legend.

## Acknowledgements

We thank John Reinitz, Claude Desplan, and Stefan Thor for antibodies; Barry Dickson, Matthieu Loius, and Vivek Jayaraman for fly stocks. Transgenic lines were generated by BestGene (Chino Hills, CA) or Genetivision (Houston, TX). Stocks obtained from the Bloomington Drosophila Stock Center (NIH P40OD018537) were used in this study. We thank Dr. Sen-Lin Lai for the immortalization fly stock, and Sen-Lin Lai, Emily Heckman, and Arnaldo Carreira-Rosario for comments on the manuscript. Funding was provided by HHMI (CQD, KML).

## Additional Information

### Funder

Howard Hughes Medical Institute

### Grant reference number

None

### Author

Chris Doe, Kristen Lee

## Figure legends

**Figure 1 – Supplement 1.**
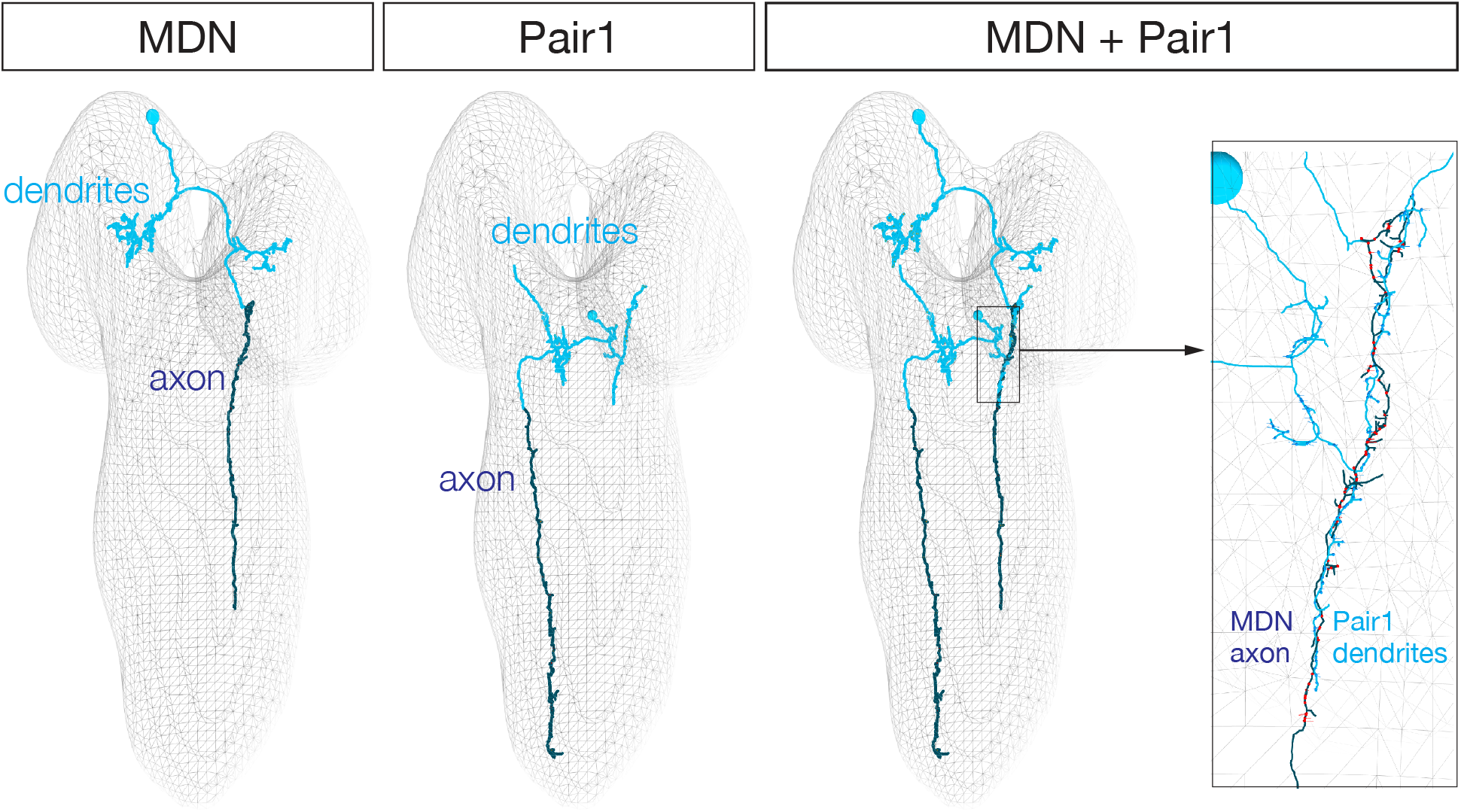
MDN axon and Pair1 dendrite target the same neuropil in the larval brain. The TEM volume of the newly hatched larva “Seymore” showing the axon (green) and dendrite (blue) domains of a single MDN and Pair1 neuron defined by the location of pre- and postsynapses. Left: MDN axon and dendrite domains. Middle: Pair1 axon and dendrite domains. Right: MDN axon and Pair1 dendrite are closely entwined the same region of neuropil (white bracket).

**Figure 3 – Supplement 1.**
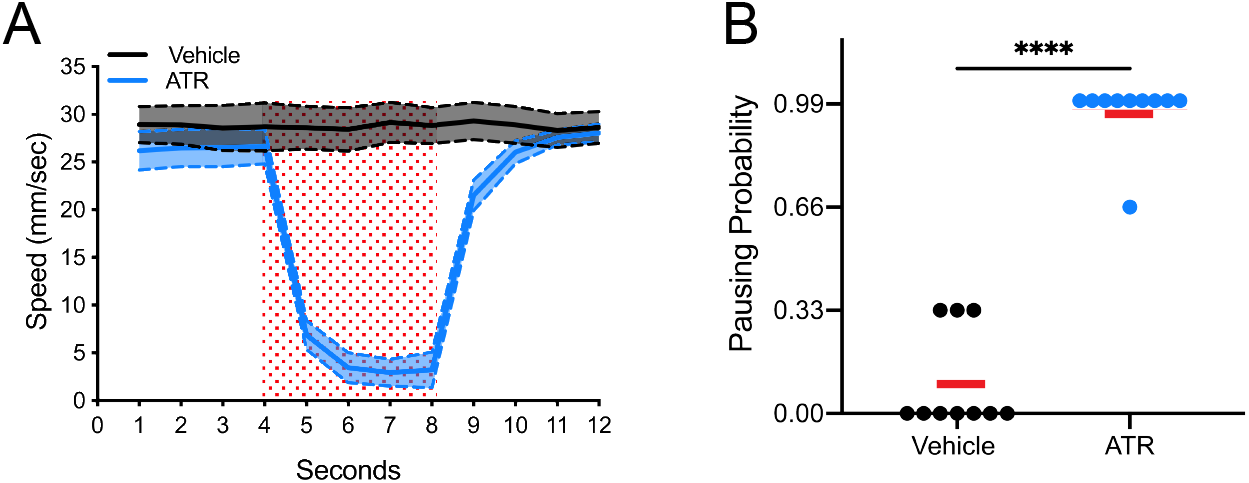
Pair1 activation for 4s arrests forward locomotion but does not cause paralysis in adults in an open field arena. (**A**) Speed (mm/sec) of adult flies in an open field arena. Flies were fed food supplemented with ATR (blue) or ethanol (vehicle, black). Red square represents the presentation of the light stimulus. (**B**) Probability of pausing upon light activation of Pair1 (ATR treatment, blue) compared to controls (vehicle treatment, black) (t-test, p < 0.001; n = 12).

